# Spatially resolved transcriptomics reveals innervation-responsive functional clusters in skeletal muscle

**DOI:** 10.1101/2022.03.31.486563

**Authors:** Chiara D’Ercole, Paolo D’Angelo, Veronica Ruggieri, Daisy Proietti, Carles Sanchez Riera, Alberto Macone, Davide Bonvissuto, Claudio Sette, Lorenzo Giordani, Luca Madaro

## Abstract

Striated muscle is a highly organized structure composed by well-defined anatomical domains with integrated but distinct assignments. So far, the lack of a direct correlation between tissue architecture and gene expression has limited our understanding of how each unit responds to physio-pathologic contexts.

Here, we show how the combined use of spatially resolved transcriptomics and immunofluorescence can bridge this gap by enabling the unbiased identification of such domains and the characterization of their response to external perturbations. Using a spatiotemporal analysis, we followed the changes in the transcriptomics profile of specific domains in muscle in a model of denervation. Furthermore, our approach allowed us to identify the spatial distribution and nerve dependence of atrophic signalling pathway and polyamine metabolism to glycolytic fibers. Indeed, we demonstrate a pronounced alteration of polyamine homeostasis upon denervation. Our dataset will serve as a resource for future studies of the mechanisms underlying skeletal muscle homeostasis and innervation.

**Graphical Abstract:** 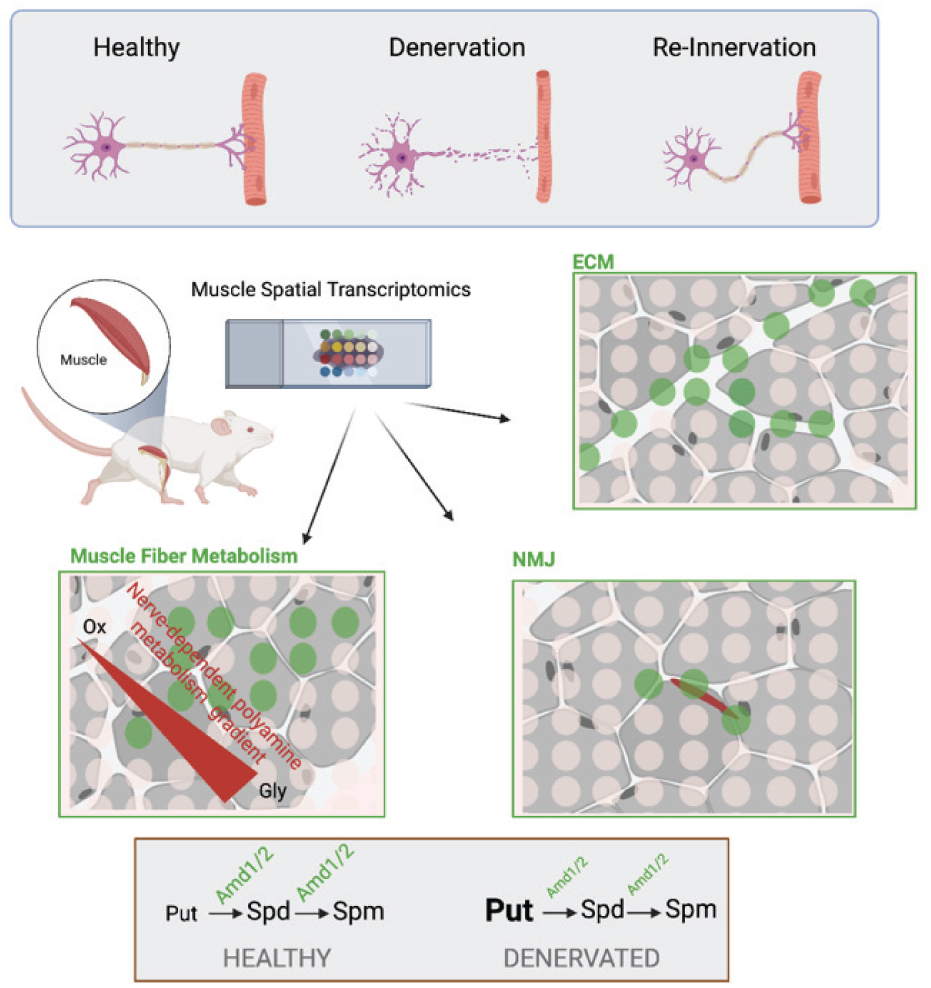

## INTRODUCTION

Skeletal muscle is a complex organ in which several mononucleated cell populations organize within specific anatomic compartments. These domains -nerves, connective, and vasculature - act altogether in coordination with muscle fibers to ensure the seamless execution of each contraction. In response to acute injury specific resident muscle populations, activate and contribute to regeneration by regulating muscle stem cell activity or cooperating with immune cells recruited from the circulatory system (Neutrophils, Macrophages) (Oprescu et al., 2020). However, muscle regeneration is not limited to myofiber repair, in order for the muscle to be functional all the different compartments need to be repaired. Indeed, voluntary contraction can only be restored through the establishment of new neuromuscular junctions (NMJs) on the newly formed myofibers (Liu and Chakkalakal, 2018). Innervation not only enables voluntary muscle control but also contributes to the maintenance of tissue homeostasis by regulating the balance between anabolic and catabolic responses(Sartori et al., 2021).

Recently, due to the advent of single-cell and single-nucleus RNA sequencing (snRNAseq), the heterogeneity of muscle cellular populations and their relative role under physiological and pathological conditions such as injury, denervation or genetic disorders have begun to be methodically surveyed (Chemello et al., 2020; Dell’Orso et al., 2019; Giordani et al., 2019; Kim et al., 2020; de Micheli et al., 2020; Nicoletti et al., 2020; Petrany et al., 2020; Petrilli et al., 2017; Proietti et al., 2021; Rubenstein et al., 2020; dos Santos et al., 2020; Schaum et al., 2018) However, these approaches have several intrinsic limitations that hinder our ability to fully understand skeletal muscle pathophysiology. In particular, most of skeletal muscle mass is composed of long, polynucleated fibers, which, due to the necessary dissociation step, are incompatible with the analysis of cytoplasmic RNAs. Moreover, functional anatomical domains such as NMJs, tendon-to-muscle connections, vessels and connective tissue can have different interactions depending on their relative proximity and locations; unfortunately, upon processing for single-cell analysis, the tissue architecture is disrupted, and this higher-order information are lost. As a consequence, the complexity of muscle tissue in pathophysiological conditions, in terms of both, cellular heterogeneity and spatial relocalization, has not yet been fully described. Despite recent advances (Lin et al., 2021; Nicoletti et al., 2020; Proietti et al., 2021), there remains an incomplete understanding regarding how traumatic events -such as nerve injury - differentially influence the specific anatomical domains and the various cellular identities in skeletal muscle. In this context, *spatial transcriptomics (ST)* could contribute to a revolution in the field of muscle “omics” as a fusion of recent sequencing technologies and classical histology(Achim et al., 2015; AL et al., 2020; Hunter et al., 2021; Rao et al., 2021).

Thus, by using ST, we aimed to characterize the discrete molecular signature of the different anatomical domains in skeletal muscle and investigate their relative role in the well characterized model of nerve crush injury. Coupling ST with immunofluorescence we first unbiasedly identified functional anatomical clusters in healthy muscle. Then, probing the muscle at different timepoints we evaluated their specific transcriptomic alterations during re-innervation. Lastly, capitalizing on ST we identified two key enzymes of the polyamine pathway whose expression is innervation-dependent and restricted to glycolytic fibers. Our data define the transcriptional profile of the different anatomical compartments in skeletal muscle and set the basis for the understanding of differential fiber sensitivity to atrophy and nerve dependent gene expression.

## RESULTS

### Spatial transcriptome analysis identifies 8 different clusters in muscle tissue

To establish a transcriptional reference of the different skeletal muscle anatomical domains and investigate their response to denervation we took advantage of the Visium Spatial Gene Expression platform (10x Genomics). Mouse limb tibialis anterior (TA, tibial muscle) and associated extensor digitorum longus (EDL) were harvested before and at different timepoints after reversible nerve injury (3 and 30 days after sciatic nerve compression - experimental workflow is shown in Figure 1A). Muscle cryosection were then placed on Visium Slide capture areas and stained. Each area contains ∼5000 RNA capture spots characterized by a specific spatial barcode that ensures that transcripts can be mapped to their original histological location. To highlight the different anatomical features of the muscle, we used anti-laminin and anti-collagen-1 antibodies, bungarotoxin (BTX) and Hoechst. This strategy enabled us to visualize fiber boundaries, extracellular matrix scaffolds, NMJs and cellular nuclei, respectively (Figure 1B and Suppl. Figure 1A and C). After imaging, tissue was permeabilized and polyadenylated RNA captured on the underlying spots. Following manufacturer protocols, we performed reverse transcription, second strand synthesis and amplification. After library preparation and sequencing, reads were processed and spatially resolved using Space Ranger software. The different datasets were then analyzed separately and integrated using Seurat 4 (Hao et al., 2021a). Lastly, data were overlaid onto the acquired image of the tissue sections (Figure 1B and Suppl. Figure 1A and C). Overall, we identified 14,417 features across 2,987 unique capture spots. Unsupervised clustering identified 8 clusters with consistent transcript abundance distribution (Supp. Figure 1E). Tissue projection showed clear spatial segregation for four out of the 8 clusters (cluster 0, 1, 3, 4) (Figure 1C - Supp. Figure 1B and D). This specific spatial arrangement readily suggested that these clusters could represent the major metabolic domains and general areas of the muscle such as different type of fibers or specific components of the matrix scaffold. Conversely, clusters 2, 5, 6, and 7 showed a sparse distribution more in line with small functional elements disperse within the tissue such as NMJ, vessel or nerves. Based on this initial observation we therefore set out to determine the identity of each of these specific cluster based on their gene expression profile (Supplementary Table 1) and their overlap with pre annotated anatomical structures in the uninjured muscle.

**Figure 1:**
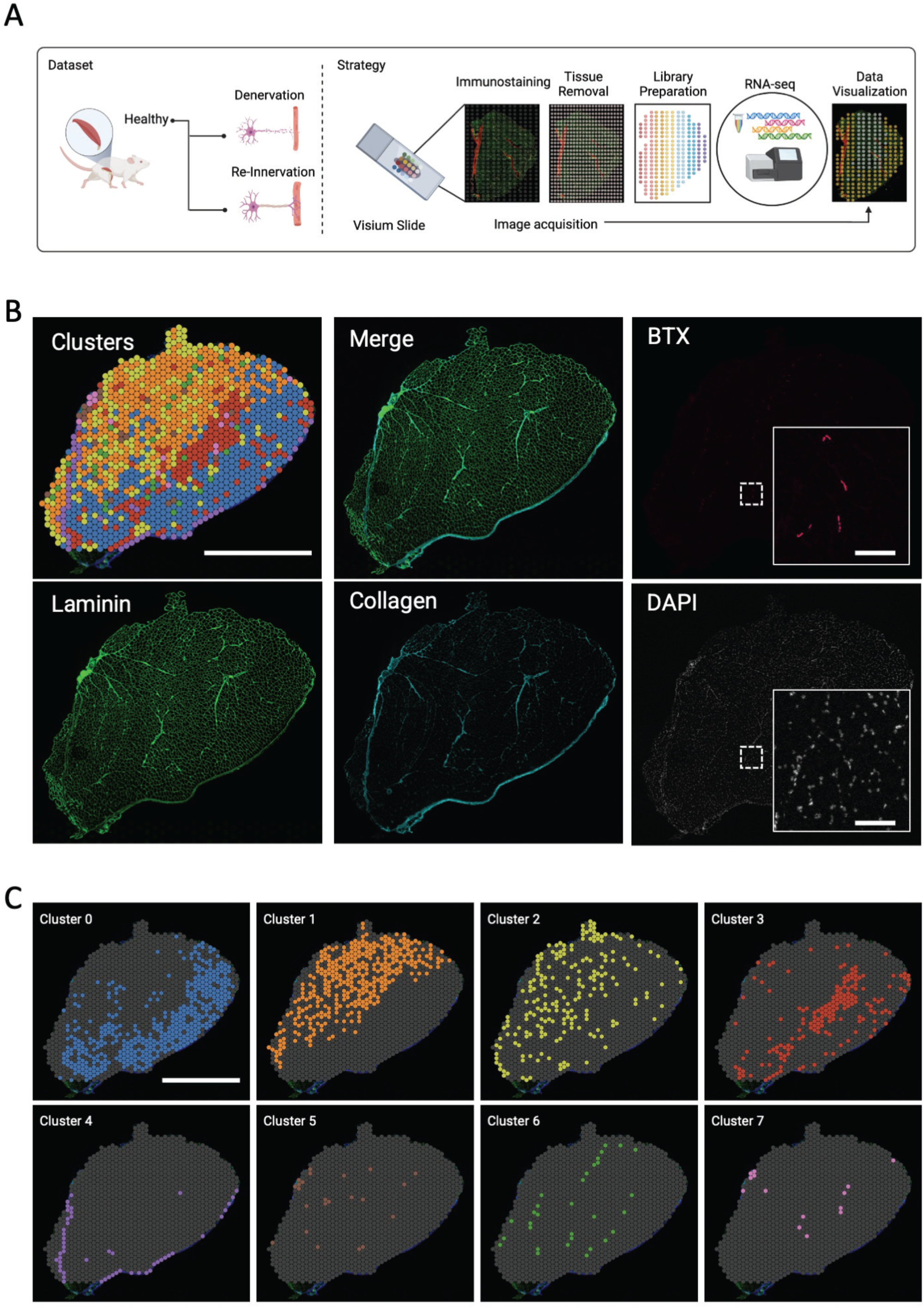
Spatially resolved transcriptomics reveals highlights discrete functional anatomical units. A) Diagram illustrating the experimental workflow for Spatial Transcriptomics (ST) analysis using 10x visium slide. B) WT mouse hindlimb cryosection (TA and EDL) used for ST analysis. Visium array spots are color-coded based on cluster assignment of the integrated dataset (Top left). Immunofluorescence staining for Bungarotoxin (top right) Laminin (bottom left) Collagen (bottom center) and Hoechst (bottom right). Scale bar, 2mm (inset Scale Bar 100um) C) Spatial distribution of each cluster. Scale bar, 2mm

### Spatial transcriptomic clustering reveals functional and structural organization of muscle tissue

We initially correlated the distribution of genes in unbiased predicted clusters with functionally characterized muscle areas. As shown by the NADH-TR staining -used to identify areas with high oxidative activity- TA muscle is characterized by a glycolytic cortex, in which muscle fibers are predominantly lightly stained, and by a darker oxidative core (Figure 2A). Indeed, staining intensity directly correlates with the number of mitochondria within a muscle fiber and reveals the characteristic pattern of finer fiber types. Based on their metabolic properties and the specific myosin subsets expressed muscle fibers can be divided in slow twitching (Type 1) and fast twitching (Type 2). Tibialis anterior is primarily composed of Type 2 fibers, which can be further subdivided into 3 main categories (2A oxidative, 2X and 2B glycolytic). Interestingly, whereas Cluster 1 was associated with the deeper oxidative core of the muscle normally characterized by the presence of 2A fibers, Cluster 0 overlapped with the glycolytic cortex of the tibialis containing mostly 2B fibers (Figure 2A). Indeed, Gene Ontology (GO) analyses of Cluster 1 genes revealed the enrichment of terms associated with mitochondrial structure and respiration (Suppl. Figure 2 Table A) and, as expected, cells in the two clusters expressed canonical markers of specific fiber types. For example, myosin heavy chain 2 (*Myh2 –* characteristic of 2A fibers), lactate dehydrogenase B (*Ldhb*) post-glycolytic enzyme and slow fiber markers such as *Myl3* were spatially restricted to Cluster 1, while the fast twitch glycolytic myosin isoform 4 (*Myh4*) and the associated fast isoform of tropomyosin (*Tpm1*) primarily cosegregated into Cluster 0 (Figure 2B). Notably, the ST profile confirmed that expression of the Mettl21c gene is confined to the tibialis and almost absent from the EDL muscle (Figure 2B), as previously reported (Wiederstein et al., 2018). Interestingly enough in Cluster1 we detected also the expression of myosin heavy chain 1, (Myh1) a hallmark of 2X fibers (Figure 2B). We speculated that, given their phenotype, intermediate between 2A and 2B in terms of contraction velocities, and mitochondrial activity (Schiaffino and Reggiani, 2011), these fibers could also be present in Cluster1.

**Figure 2:**
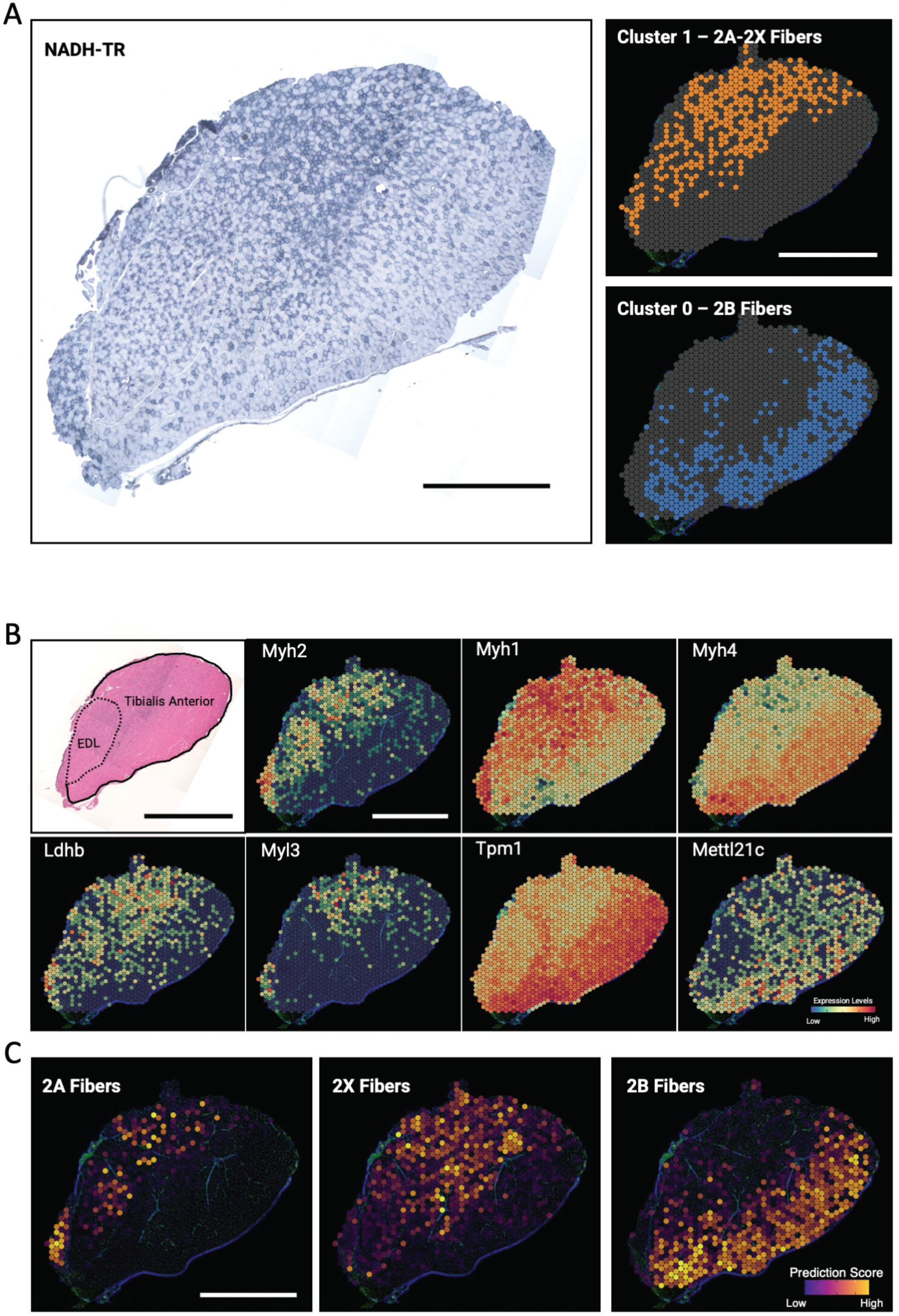
Cluster 0 and Cluster 1 correspond to Glycolitc and Oxydative Fibers. A) NADH–TR staining of consecutive section (Scalebar 1mm); Cluster 0 and Cluster 1 pattern display an overlap with NADH-TR low and NADH-TR high expressing fibers. Scale bar 2 mm B) Serial section stained with Hematoxylin and Eosin illustrating muscle localization (top left) and relative expression levels of fiber-type specific marker genes from the ST data projected over the tissue space. Scale bar 2 mm C) Projection on the tissue space of the prediction Score for IIa, IIx and IIb fibers based on snRNAseq from WT mouse (Chemello et al.) integrated with ST dataset. Scale bar 2 mm

To test this hypothesis and further validate our annotations, we integrated our ST data with a previously published single-nucleus RNA sequencing (WT dataset from Chemello et al., 2020), and we spatially mapped cell type predictions for the different fiber types (see Methods). Data intersection confirmed that fiber type metabolism was correctly predicted based on the spatial distribution of the myosin types identified in each cluster (Figure 2C). While the glycolytic portion, overlapping with Cluster0, was predicted to contain type 2B fibers (fast glycolytic), the portion overlapping with Cluster1 consisted of type 2A fibers (fast oxidative), and type 2X fibers, which are less glycolytic than type 2B fibers (Figure 2C).

Next, we analyzed the molecular signature of the other two clusters that presented a clear spatial distribution, Cluster3 and Cluster4. Upon GO analysis, both Clusters 3 and 4 displayed an enrichment in ECM components (Suppl. Figure 3 Table A). However, while Cluster 3 overlapped considerably with the intramuscular matrix scaffold, Cluster 4 clearly identified the epimysium surrounding the muscle (Figure 3A and Suppl. Figure 3B). Almost perfect overlap was obtained between the localization of Cluster4 and the localization of collagen 1 surrounding the muscle (validated by Immunofluorescence and Sirius red staining - Figure 3A - inset). As expected, the genes enriched in this cluster are extracellular matrix components such as collagen subunits (Col12a1 and Col1a1) and matrix-associated molecules (Fmod and Timp2) (Figure 3B).

**Figure 3:**
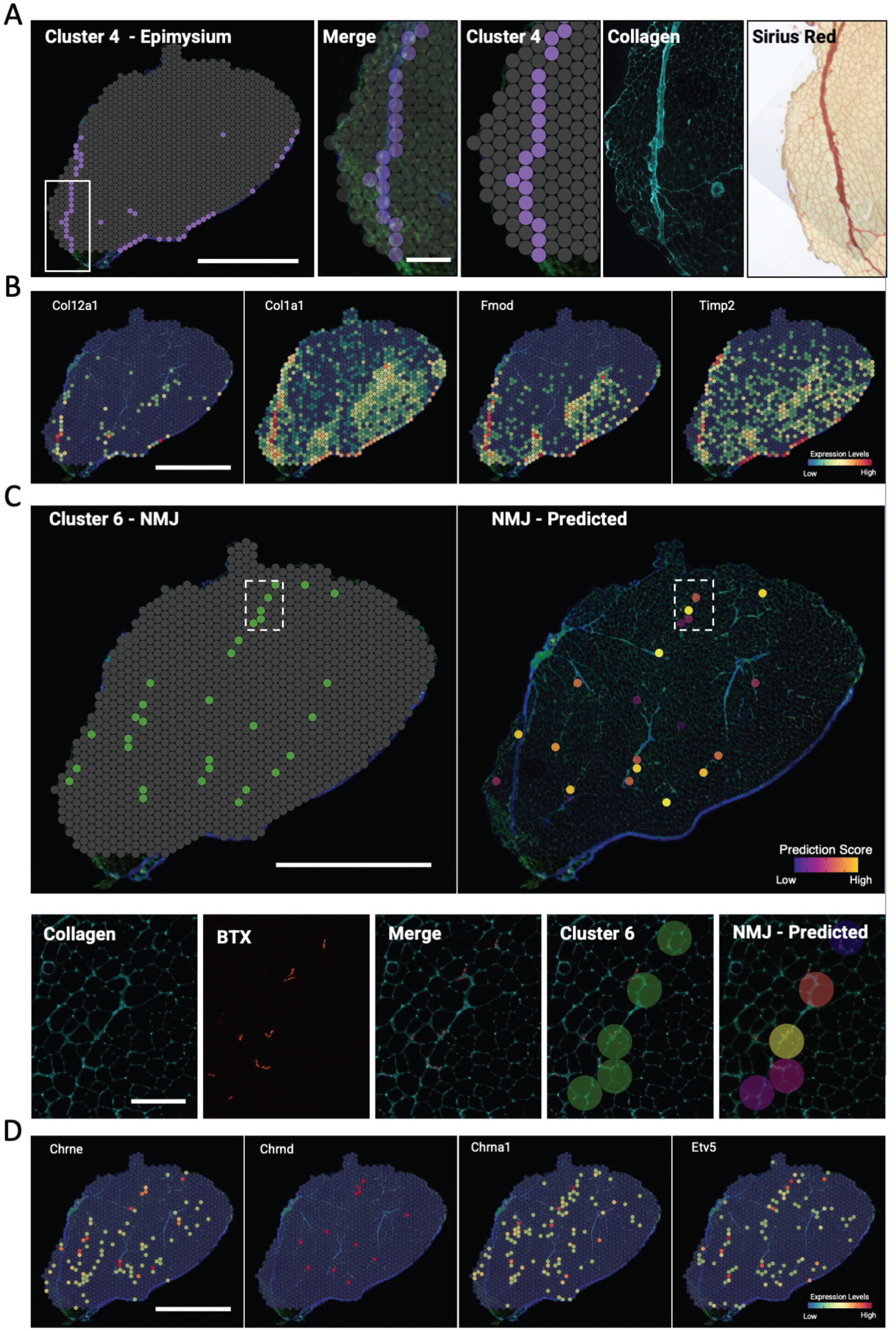
Cluster 4 and Cluster 6 correspond to muscle perimysium and neuromuscular junction. A) Spatial distribution of cluster 4(left) Scalebar 2mm. Overlay of cluster 4 distribution over tissue space, collagen staining of the region highlighted and Sirius red Staining of serial section. Scalebar 100um B) Relative expression levels of genes enriched in Cluster 4. Scalebar 2mm C) Spatial distribution of cluster 6 (top left) and projection of prediction score for NMJ cells (top right). Scalebar 2mm. Enlargement of the region highlighted in the white box (bottom), Immunostaining for collagen and bungarotoxin respectively (left) and overlay of cluster 6 and NMJ prediction score (right). Scalebar 100um D) Relative expression levels of NMJ specific marker genes from the ST data projected over the tissue space. Scale 2mm

Given the complexity of muscle tissue, we wondered whether ST analysis might not only reveal more abundant clusters related to muscle fibers but also identify more restricted morphofunctional zones specific to this tissue. We therefore analyzed the gene expression signature of the sparse clusters (Cluster2, Cluster 5, Cluster6, Cluster7) and their relative spatial proximity to known morphological structures. Upon visual inspection, Cluster 2 did not present a recognizable distribution pattern nor overlap with clearly detectable anatomical compartments. Furthermore, we could not detect a strong gene expression signature for the cluster. However, it must be considered that each spot contains data from an ensemble of different cell types, and it is possible that, transition areas where such mix is more diverse could be clustered together. This could explain the weak gene signature and the associated generic GO terms (muscle contraction and muscle system process – Suppl. Figure 3C). Additionally, it is worth mentioning that the expressed genes identified included Alas2 and different hemoglobin transcripts, suggesting specific erythroid enrichment. Thus, at least some of these regions could contain small blood vessels.

Cluster 6 spots, on the other hand, nicely colocalized with acetylcholine receptor across muscle sections (stained with Alexa 594-conjugated BTX). In line with the spatial pattern GO analysis of genes enriched in of Cluster6 contained terms associated with NMJs and synapses (Suppl Figure 3 Table C). As further validation, NMJ location prediction based on snRNA-seq integration overlapped with Cluster 6 spots (Figure 3C). Lastly, representative genes of this cluster encode acetylcholine receptor subunits (Chrne, Chrnd, Chrna1) or transcription factors known to be expressed in subsynaptic nuclei (Etv5)(Kim et al., 2020) (Figure 3D).

Interestingly enough, also Cluster5 and Cluster7 correlated with histological structures, associating respectively with nerves and large vessels. Indeed, Cluster5 was enriched in glial (Plp1) and myelin (Pmp22, Mpz, Mbp) associated factors, while Cluster7 expressed endothelial or smooth muscle-associated genes (CD31-Pecam1, Vwf, Myh11 and Acta2) (Figure 4A to C).

**Figure 4:**
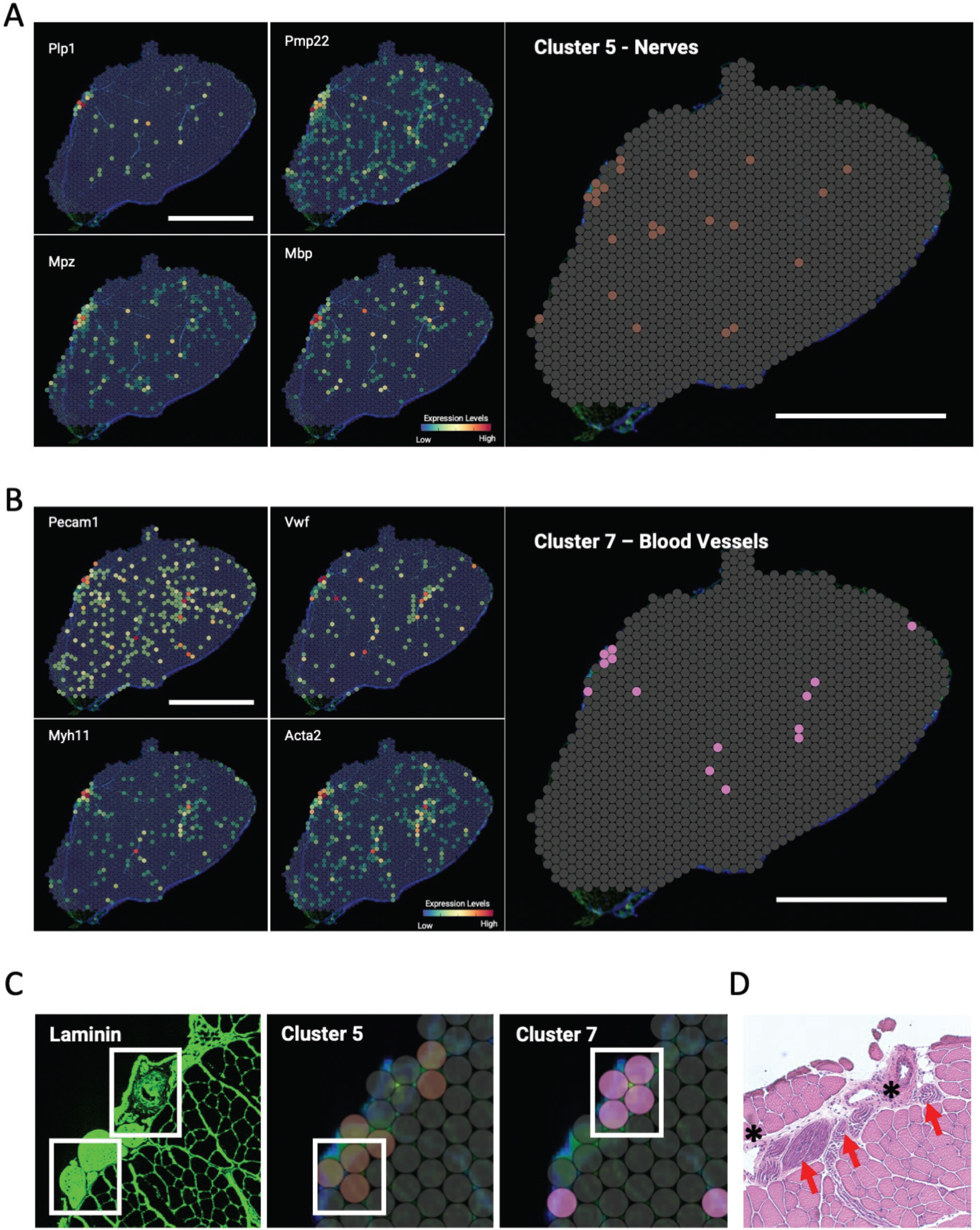
Cluster 5 and Cluster 7 correspond to Nerves and Blood Vessels respectively. A) Relative expression levels of genes enriched in Cluster 5. Spatial distribution of cluster 5(right). Scalebar 2mm B) Relative expression levels of genes enriched in Cluster 7. Spatial distribution of cluster 7(right). Scalebar 2mm C) Laminin staining of a specific region of the muscle where nerve and vessels are in close proximity. White Boxes highlight the relative distribution of Cluster 5 and Cluster 7 and the overlap between the anatomical structure and the transcriptomic unit. Scalebar 100um D) Serial section stained with Hematoxylin and Eosin illustrating nerve localization (red arrows) and vessel localization (asterisk). Scale bar 100 um

Notably, spatial gene expression data allowed the resolution of structures that are closely associated with each other anatomically. For example, Clusters 5 and 7 discriminated structures attributable to nerves and vessels, respectively, even when they were located in close proximity (Figure 4C-D). Taken together this data suggest that, despite the relatively “large” diameter of the capturing spot (55μm) this strategy can be used to precisely identify the different morpho-functional domains in muscle.

### Spatial transcriptome clustering analysis of transiently denervated muscle

Once correctly annotated the different clusters in the unperturbed muscle we proceeded to investigate their response to reversible denervation. Sciatic nerve compression is characterized by a stereotypical induction of the atrophic response that is resolved upon reinnervation (Magill et al., 2007). Complete denervation of TA and EDL is observed three days after injury (Suppl. Figure 4A) with the consequent activation of the muscle-specific ubiquitin ligases Atrogin-1 (Fbxo32) and MuRF1 (Trim63) in both muscles (Figure Supp.4B). Eventually both ubiquitin ligases will be downregulated upon progressive repair and reinnervation around day 30 (Suppl. Figure4B). Our bulk ST data recapitulated the main features of the model with an enrichment of the Foxo signaling pathway 3 days after nerve injury in a pseudobulk analysis in conjunction with a peak in expression profile of Atrogin-1 and MuRF1 (Suppl. Figure 4 C and D and Suppl. Table 2). We then analyzed the specific gene expression profile of each cluster at 3 days and 30 days after denervation (Figure 5A and Suppl. Table 3). Similarly, we identified the upregulation of genes related to atrophy in specific clusters such as the growth arrest and DNA damage-inducible 45α (Gadd45a), the ubiquitin ligase Itch (Nerves) and the homeobox gene Tgif (Blood Vessels) or autophagy related genes Beclin1 (Becn1) and Gabarapl1 in both Nerve and Blood Vessel Clusters (Figure 5B). Interestingly, we noticed that the induction Atrogin-1 and Murf1 was primarily localized in the glycolytic cortex of the TA muscle (Figure 5C compared with Figure 2A). In addition, 2B cluster displayed higher transcripts levels when compared to the 2A-2X cluster (figure 5D). This observation suggests that the distribution pattern of the atrophic response depends on the muscle fiber composition, consistent with data in the literature indicating that glycolytic muscle fibers are more susceptible to muscle atrophy than oxidative muscle fibers (de Theije et al., 2015; Wang and Pessin, 2013).

**Figure 5:**
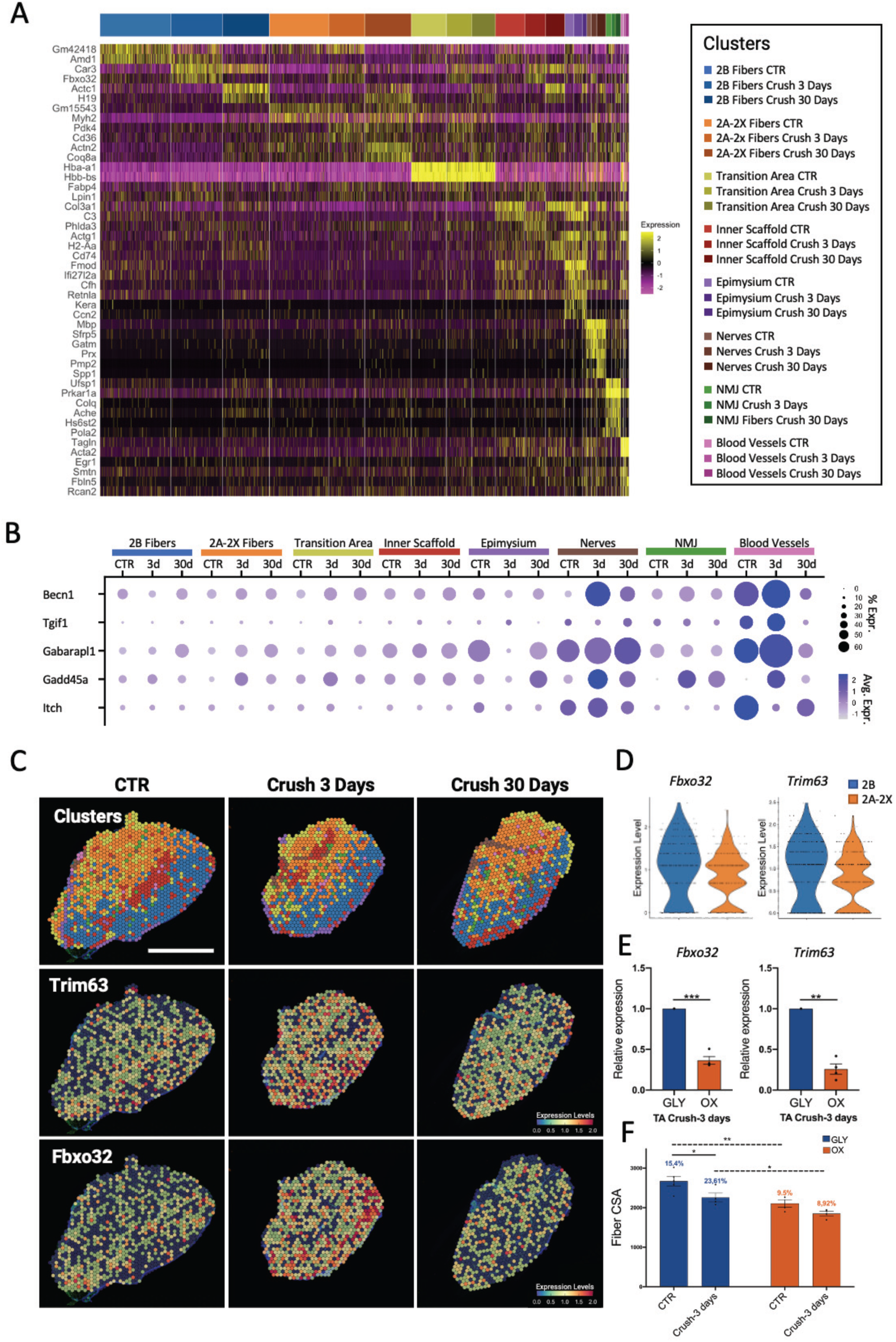
ST reveals discrete dynamics during denervation-induced atrophy. A) Heatmap showing expression values of the 2 most variable genes for each cluster in each timepoint after differential gene expression analysis B) Dotplot illustrating the relative expression and distribution of the cluster-specific atrophy genes. C) Cluster distribution and relative expression levels of Fbox32 Trim63 transcripts during denervation- reinnervation process. Scale 2mm D) Violin plot showing the expression of Fbox32 Trim63 in glycolytic and oxidative fibers clusters. E) Relative expression of Fbox32 Trim63 measured after laser microdissection of TA 3 days after denervation. (n = 4, values represent mean SD, **P < 0.01, ***P < 0.001, ; by 2-tailed Student’s t test). F) Mean CSA of muscle fibers (n = 4, values represent mean SD, *P < 0.05, ****P < 0.01, by 1-way ANOVA Tukey’s multiple-comparison test).

We confirmed this trend in our observations, via qPCR analysis of laser-microdissected glycolytic (Gly) or oxidative (Ox) areas of the muscle sections (see scheme in Suppl. Figure 4E). MuRF1 and Atrogin-1 were more highly expressed in the glycolytic cortex of the denervated TA muscle than in the oxidative core (Figure 5E). Interestingly, although the reduction in fiber area was rather limited at 3 days of denervation, we observed by separately measuring the fibers in the glycolytic cortex and those in the oxidative core (strategy shown in Suppl. Figure 4E) that the reduction was significantly more pronounced in the glycolytic portion (Figure 5F), where atrogenes (Atrogin-1 and Murf1) were more highly expressed.

### Nerve-dependent polyamine synthesis-related enzyme pattern across muscle

Obtaining a detailed map of the spatial localization of specific genes in muscle could be pivotal for pinpointing new signaling pathways involved in the pathophysiology of the tissue. To this end, the possibility of identifying the specific distribution of certain pathways in functionally different areas of muscle will be of particular interest. As a proof of concept, we identified the spatial distribution of the genes encoding Smox and Amd1, enzymes related to polyamine (PA) synthesis (Figure 6A – highlighted in bold), which showed a pattern reminiscent of the distribution of glycolytic fibers in the TA (Figure 6B). As an initial finding, ST data revealed in the CTR a greater expression of these genes in the TA than in the EDL (Figure 6B), which was confirmed by qPCR analysis (Figure 6C). Specifically, Amd1, Amd2, Smox, and Sms expression profile was largely overlapping the glycolytic cortex of the muscle (Figure 6B and Supp-Figure 5A) with a clear enrichment in the 2B Fiber cluster when compared to 2A-2X cluster (Figure 6D).

**Figure 6:**
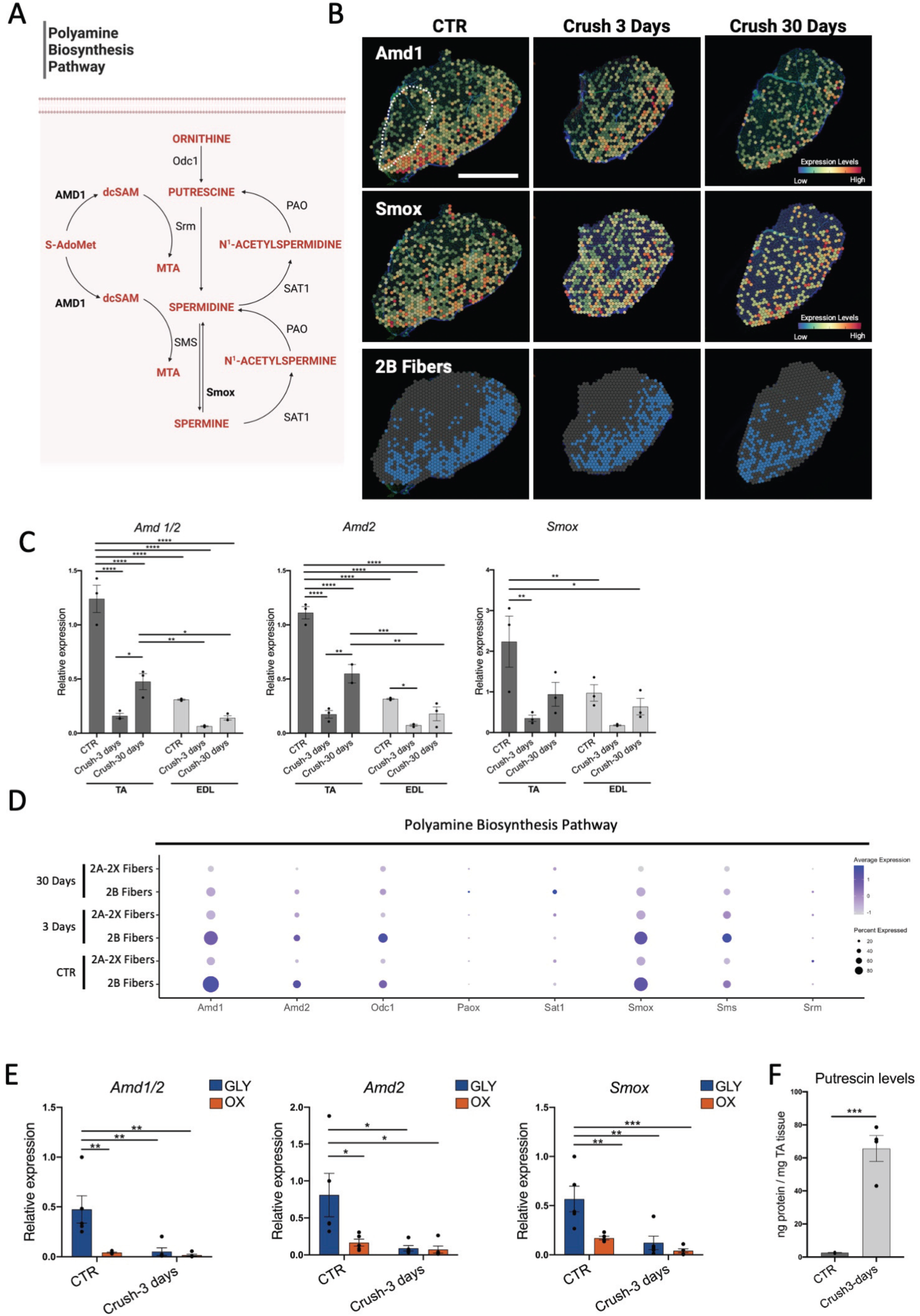
ST identifies innervation dependent pathways. A) Diagram Illustrating Polyamine Pathway B) Cluster 0 distribution and relative expression levels of spermidine pathway component (Amd1 and Smox) scale 2mm (dotted line highlights EDL muscle) C) Relative expression of Amd1/2, Amd2 and Smox during denervation in TA and EDL muscle (n = 3, values represent mean SD, *P < 0.05, **P < 0.01,***P < 0.001, ****P < 0.0001; by 1-way ANOVA Tukey’s multiple-comparison test). D) Dotplot illustrating the relative expression and distribution of the Spermindine pathway. E) Relative expression of Amd1/2, Amd2 and Smox after laser microdissection of TA 3 days after denervation. (n = 4, values represent mean SD, *P < 0.05, **P < 0.01,***P < 0.001; by 1-way ANOVA Tukey’s multiple-comparison test). F) GC-MS quantitative analysis of Putrescine level in control (CTR) and denervated (Crush 3 days) muscles. (n = 4, values represent mean SD,***P < 0.001 by 2-tailed Student’s t-student test).

It is noteworthy that, concomitantly with a reduction in overall expression in the glycolytic cluster, we observed a partial loss of the spatial restriction of these genes (Figure 6B and D). Intriguingly, at 30 days after the induction of nerve damage-when innervation is restored- Amd1, Amd2 and Smox expression in the muscle is still decreased compared to CTR levels (Figure 6C). This could suggest that more time or more extensive maturation of NMJs is needed to re-establish the correct expression of these markers.

We validated this observation through laser microdissection; indeed, while the expression of Amd1, Amd2 and Smox was high in the glycolytic portion of the TA in control tissue, after denervation, due to their reduced expression in glycolytic fibers, the relative levels of these genes were similar between the glycolytic and oxidative fractions of the muscle (Figure 6E). We speculated that such change in expression levels could result in an imbalance in the PA pathway. The decarboxylation of S-adenosylmethionine (SAM), catalyzed by Amd1/Amd2, produces an aminopropyl group that acts as a substrate together with Putrescine for the Spermidine synthase and subsequently to form Spermine (Figure 6A). Thus, reduction of Amd1/Amd2 expression may results in Putrescine accumulation. Indeed, as shown by gas chromatography-mass spectrometry (GC-MS) quantitative analysis, we observed a dramatic change in polyamine ratio 3 days after denervation (Figure 6F). A mean 25-fold increase of Putrescine was revealed as results of Amd1 limitation. However, although their precursor is significantly increased the amount of spermidine and spermine does not change significantly (Figure Suppl. 5C). Intriguingly, in support of this observation an induction of Odc1, required to synthesize Putrescin from Ornithine, is upregulated in denervated muscle (Suppl. Figure 5D).

Collectively, these data show how ST could be used to identify variation in the spatial patterning of specific genes and pathways in skeletal muscle – as in example occurs to the PA pathway as result of denervation.

## DISCUSSION

Here we applied Spatial Transcriptomic Visium assay coupled with immunofluorescence to profile the transcriptional signature of distinct functional histological domains in skeletal muscle. Using a spatiotemporal analysis, we characterized their transcriptional response in a model of reversible nerve damage. Skeletal muscle is a complex organ in which the main motor unit, the fiber, is tightly interconnected with structures of different origins and heterogeneous cellular composition. Recent advances in terms of single-cell RNA sequencing have given new input to the field, enabling the unbiased transcriptomic analysis of all tissue-resident populations. However, due to the lack of positional information current scRNA-seq and snRNAseq methods are still unable to fully render the complexity of the different anatomical domains in muscle, ultimately providing a list of cellular components but limited insights into the coordinated activity of the whole functional unit. On the contrary, by contextualizing gene expression within its true location, ST could provide detailed information on how different structural units acts and synergize.

Indeed, our data faithfully recapitulate all the molecular events triggered by reversible nerve loss and offer a detailed temporal snapshot of the changes that occur in the different functional domains. In line with what has been extensively described in literature we readily detected the activation of the Foxo-mediated atrophic mechanism leading to reduced muscle mass. Furthermore, our analysis highlighted the differential activation of atrophy-related genes in specific anatomical districts. In particular we showed that Beclin1 and Gabarapl1 are upregulated in both Nerves and Vessels thus suggesting a predominant early activation of autophagy in those specific clusters. In addition, we confirmed differences in the extent of the atrophic response in oxidative vs. glycolytic fibers, with earlier and more extensive activation of Murf and Atrogin-1 in the latter. If fiber types indeed activate distinct but consistent transcriptomic programs, differences in the atrophic response that have thus far been considered intermuscular differences would thus likely be dependent only on the fiber type composition (Brocca et al., 2017); however, it could conversely and more intriguingly be speculated that each muscle could trigger a tailored atrophic program that is then coordinately executed by the different fiber types acting interdependently. Indeed, our data support the hypothesis that within the same muscle the specification of fibers results in a different susceptibility to external stimuli by activating with various intensity an atrophic program in distinct anatomical zones of the same muscle.

As a proof of concept that this technology can be successfully applied to skeletal muscle to identify and investigate the spatial distributions of specific signaling pathways within the tissue, we also showed how the expression of enzymes of the polyamine synthesis pathway follows a fiber metabolic gradient. In fact, genes encoding for key enzymes of this pathway tend to be more highly expressed in the glycolytic portion of the TA muscle than in the oxidative core. The full elucidation of the role of this signaling pathway in different fiber types will require further studies. It is nevertheless worth mentioning that another group recently demonstrated that PA metabolism is required for correct Drosophila locomotor function (Coni et al., 2021). Furthermore, the spatial regulation of gene expression seems to be directly mediated by local innervation, since after nerve injury, we observed a loss of this gradient as a result of reduced expression of these genes in the glycolytic compartment. Moreover, the reduction of key enzymes for the conversion of putrescine into spermidine and then into spermine has the effect of dramatically accumulating putrescine in denervated muscle. In fact, the ratio between the different polyamines in the denervated muscle is upset resulting in a massive accumulation of putrescine. This could be directly involved in the atrophic phenotype since while spermidine as been shown to prevent atrophy and neurodegeneration (Clarkson et al., 2004; Noro et al., 2015b, 2015a; Sharma et al., 2018), putrescine has been associated with neurodegenerative phenomena (Camón et al., 2001; Plewa et al., 2021; Virgili et al., 2006). Although the result of the altered ratio of polyamines in denervated muscle requires specific studies, this observation supports the idea that innervation is required not only to support motor function, but also to regulate the gene expression profile of innervated fibers and more importantly to govern skeletal muscle functional cluster determination by driving compartmentalized transcriptional regulation.

These events are therefore related to a distinct abundance in the expression of factors that are then prevalent in determining areas of the muscle both in basal conditions (polyamine metabolism) and after alteration of homeostasis (atrogenes). In a translational perspective, this could lead to a better understanding of how some muscle fibers are more resistant to atrophy and thus promote a shift towards this type of fiber in a therapeutic perspective.

Here, we only focused on TA and EDL muscle groups alone which are almost exclusively composed of fast type fibers. In the future it would be interesting to use TS to analyze atrophy in different muscle groups with a more divers fiber composition to assess whether the same fiber types activate the same transcriptomic program in different muscles and to highlight differences with slow twitch muscle fibers (e.g. predominant in Soleus muscle). In addition, exploring the regeneration process at later time after denervation would be useful to more accurately define the molecular events that occur after nerve injury. Finally, parallel analysis of muscles undergoing nonreversible denervation would help to understand which signals are necessary to drive muscle remodeling and reinnervation and which are detrimental in mediating the catastrophic effects of innervation loss.

Recently, another spatially resolved skeletal muscle dataset has been published (McKellar et al., 2021). This dataset clearly showed the potential of a pairing spatially resolved transcriptomics with scRNA-seq to investigate cell-cell interactions. Yet the use of Hematoxylin Eosin staining as a spatial reference and the lack of a reference muscle in basal condition limited the detailed analysis of specific morphological structures. With our approach we have achieved molecular-level discrimination of areas within the muscle with differences in metabolic activity as well as functional aggregations of finer-scale areas, such as clusters of genes restricted to the extracellular matrix, vessels, nerves, and even muscle junctions. This enabled us to precisely detail the transcriptional signature within those anatomical clusters that would otherwise have been hidden by bulk RNA-seq or scRNA-seq. However, resolution remains a main hurdle to the potential applications of this technology, as each spot ideally contains one to ten cells. Several in silico methods have been developed to deconvolute or infer single cell information from a single spot (Andersson et al., 2020; Cable et al., 2021; Dong and Yuan, 2021; Elosua-Bayes et al., 2021; Hao et al., 2021b; Sun et al., 2022; Wang et al., 2019; Zhao et al., 2021) nonetheless these approaches often require scRNA-seq or bulk RNA-seq data as initial references. Eventually, the advent of more dense spot arrays will be overcome this technical bottleneck. In muscle, the limit in resolution is partially compensated by the large size of fibers that make less likely the colocalization of different fibers within a single spot. Despite the current limitations we foresee that the use of ST technology will be instrumental to understand the molecular dynamics occurring at the local level, such as stem cell-niche interactions and, by clarifying where a particular event occurs in terms of tissue localization, will contribute to elucidate the response of specific histological functional units to external perturbations such as traumatic injury or chronic inflammatory conditions.

In summary, our study provides a detailed molecular profiling that will serve as a steppingstone to investigate the mechanism underpinning muscle specific response to denervation at morpho-functional level -as in the case of differential fiber sensitivity to atrophy –.Furthermore, by providing a spatially resolved transcriptomic reference of the unperturbed muscle will allow the identification of new cellular or structural interactions in response to any perturbation of the homeostatic balance.

## ACKNOWLEDGMENTS

This work was supported by Roche per la Ricerca 2019 and AFM Telethon starting grant #23075 and Istituto Pasteur - Fondazione Cenci Bolognetti (to LM); The authors would also like to thank E. Aleo, Institute of Applied Genomics, Udine, Italy, for NGS-Seq library preparation and sequencing, PL. Puri and Bouche M. for helpful advice in drafting the manuscript.

## AUTHORS CONTRIBUTIONS

M.L and G.L designed experiments and analyzed results. D.C., D.P., R.V., P.D., C.S.R., performed and analyzed experiments. B.D and S.C. design and perform microdissection experiments. A.M. perform polyamine quantification experiments. M.L. and G.L wrote the manuscript.

## Methods

### Resource availability

#### Lead contact

Further information and requests for resources and reagents should be directed to and will be fulfilled by the lead contact, Madaro Luca

#### Materials availability

The reagent and material for this study will be available on reasonable request.

#### Data and code availability

All data supporting this study are available within the Supplementary Information file or from the corresponding author upon reasonable request.

### Denervation

Unilateral hindlimb denervation was performed by clamping the left sciatic nerve under anesthesia via intraperitoneal injection of 40 mg/kg ketamine (Zoletil®) and 10 mg/kg xylazine (Rampum). After the exposure of sciatic nerve, cutting the skin near the knee, the nerve was crushed with chirurgical tweezer three times for 10 sec each time. The lesion was sutured with 3M Vetbond Tissue Adhesive after the operation. After surgery, the mice were monitored until the sacrifice date. After the operation the animals were monitored and the actual denervation assessed by a grip test.

### Spatial RNA sequencing library preparation

TA muscles were isolated, embedded in OCT (VWR), and flash frozen in liquid nitrogen for 15 seconds, after those muscles were stored at temperature of -80 until the cryosection. Spatially tagged cDNA libraries were built using a Visium Spatial Gene Expression 3' Library Construction v1 Kit (10x Genomics, PN-1000187; Pleasanton, CA). The optimal tissue permeabilization time for 10 μm-thick sections was found to be 15 min using a 10x Genomics Visium Tissue Optimization Kit (PN-1000193). Immunofluorescence-stained tissue sections were imaged using a Zeiss confocal microscope (LSM 900 with Airyscan2) with a mechanical plate, and the images were then edited using ImageJ® software. The primary probes used for immunofluorescence were rabbit anti-laminin (1:200), mouse anti-Col1a (Col-1) (1:200), and BTX 488 (1:200). Antibody binding specificity was revealed using secondary antibodies coupled to Alexa Fluor 594 or 647. Libraries were generated from cDNA following the manufacturer’s instructions and checked with both a Qubit 2.0 Fluorometer (Invitrogen, Carlsbad, CA) and an Agilent Bioanalyzer DNA assay (Agilent Technologies, Santa Clara, CA). Libraries were then sequenced in paired-end 150 bp mode on a NovaSeq 6000 system (Illumina, San Diego, CA). The frames around the capture area on the Visium slide were aligned manually, and spots distributed across the tissue were selected using Loop Browser v5.0.0 software (10x Genomics). The sequencing data were then aligned to the mouse reference genome (mm10) using the Space Ranger v1.0.0 pipeline (10x Genomics) to generate a feature-by-spot-barcode expression matrix. Analysis was performed using Seurat v4 (the R workflow can be assessed and reproduced in R Markdown - see data and code availability). Briefly, each section was loaded using the Load10X_Spatial () function and preprocessed separately by applying sctransform normalization. After this initial step, sections were integrated using IntegrateData () with the normalization method parameter set to “SCT” and using all features from the different datasets as features for integration. Integrated dataset was normalized using SCT. After clustering and cell population identification, the most highly differentially expressed genes were identified using the ‘‘FindAllMarkers’’ function with the following parameters: only.pos = TRUE, min.pct = 0.25, and logfc.threshold = 0.25. Cell type prediction scores were inferred using the publicly available snRNA-seq WT dataset from Chemello et al. as a reference (Chemello et al., 2020). Anchors between the two datasets were identified using FindTransferAnchor with the following parameters (normalization.method=“SCT”,= and reduction = “cca”), and label transfer was performed with the default parameters. GO enrichment analysis was performed with gprofiler2 using as input cluster markers present at least in 50% of each cluster (pct.1>0.5). Pseudobulk data were obatained using “AverageExrepssion” functions and subsequently filtered for expression above 0.1 and lg2FC (3days vs CTR) >-0.3 or < 0.3. KEGG pathway analysis of genes upregulated in 3days vs CTR was performed using Enrichr (Kuleshov et al., 2016). After analysis, for visualization purposes, images were reoriented to display the EDL at the bottom-left corner.

### Laser Microdissection

TA muscles were isolated, embedded in OCT, flash frozen in liquid nitrogen and stored at -80 °C. Ten-millimeter frozen sections cut on a cryostat (Leica CM1850) were mounted on PET-membrane 1.4 mm frame slides (Leica) previously cleaned with RNase Away (Molecular Bio Products) and UV treated for 45 min under a sterile hood. NADH-TR stained muscle (protocol of staining described below) was performed to visualize the tissue structure. The glycolytic and oxidative portions of the TA muscles of 5 mice in the CTR group and 5 mice in the Crush - 3 days group were microdissected with a laser microdissection system (Leica LMD6) and recovered in RNAlater reagent (QIAGEN).

### RNA analysis by quantitative PCR

Total RNA was extracted from total muscle or dissected specimens (look above) using Qiagen RNeasy mini kits following the manufacturer’s protocol. Total RNA was quantified with a NanoDrop ONE^c^ spectrophotometer (Thermo Scientific). First-strand cDNA was synthesized from the total RNA using a PrimeScript™ RT Reagent Kit with gDNA Eraser (Takara) following the manufacturer’s protocols. The generated cDNA was used as a template in real-time PCR with Fast Q-PCR Master Mix (SMOBIO), which was run on a QuantStudio 7 Flex (Applied Biosystems, Thermo Scientific) machine with three-step amplification (1. template denature and enzyme activation step 95°C 2 min for 1 cicle – 2. denature step 95°C 15 sec and 3. anneling/extension steps 60°C 60 sec, 2 and 3 steps for 40 cycles) and melting curve analysis. Each reaction in the quantitative real-time PCR consisted of 2x Fast Q-PCR Master Mix (SYBR, ROX), 2.5 mM forward and reverse primers and 10 ng of cDNA. Relative gene expression values were normalized by dividing the specific expression value by the glyceraldehyde 3-phosphate (GAPDH) or actin beta (Actb) expression value and calculated using the 2^−ΔΔCT^ method.

### Histological staining

For Sirius red staining muscle cryosections were fixed for 1 h at 56 °C in Bouin’s solution and then stained in a Picro-Sirius red (0.1%) solution for 1 h protected from light. After a brief wash in acidified water (0.5% vol/vol), the sections were fixed in 100% ethanol, and the final dehydration step was performed in 100% toluene. The sections were mounted with EUKITT mounting medium (Sigma– Aldrich) and visualized using a Zeiss Imager.A2.

For H&E staining, the sections were fixed in 4% PFA for 10 min, washed in PBS and then stained in hematoxylin for 12 min and eosin for 30 sec. For NADH staining, the sections were placed in buffer solution (0.1 M Tris HCl, pH 7.5, in ddH_2_O) for 5 min. Then, they were stained with NADH solution (2 mg of NADH and 4 mg of NBT in 0.1 M Tris HCl, pH 7.5, in ddH_2_O) for 1 h at 37° in the dark. After removing the dye, all muscle sections were further dried in gradually increasing concentrations of ethanol in water, and after fixation in 100% toluene, they were mounted with EUKITT mounting medium (Sigma–Aldrich). The sections were imaged using a Zeiss Imager A2.

### Polyamine analysis

Polyamine content was determined by gas chromatography-mass spectrometry (GC-MS) and the values were normalized by muscle weight. TA muscles pulverized in liquid nitrogen were resuspended in 0.2 M HClO and homogenized in an ice-bath using an ultra-turrax T8 blender. The homogenized tissue was centrifuged at 13,000× g for 15 min at 4°C; 0.5 mL of supernatant was spiked with internal standard 1,6-diaminohexane and adjusted to pH≥12 with 0.5 mL of 5 M NaOH. The samples were then subjected to sequential N-ethoxycarbonylation and N-pentafluoropropionylation. For DM2 samples, biopsies were also resuspended in 0.2 M HClO4 and processed as described above. GC-MS analyses were performed with an Agilent 7890B gas chromatograph coupled to a 5977B quadrupole mass selective detector (Agilent Technologies, Palo Alto, CA). Chromatographic separations were carried out with an Agilent HP-5ms fused-silica capillary column. Mass spectrometric analysis was performed simultaneously in TIC (mass range scan from m/z 50 to 800 at a rate of 0.42 scans s–1) and SIM mode (put, m/z 405; spd, m/z 580, N1-acetyl-spm, m/z 637; spm, m/z 709).

### Figure design

The graphical abstract, Figure 1A, and Figure 5A were created with BioRender (https://biorender.com/)

## QUANTIFICATION AND STATISTICAL ANALYSIS

### Statistical analysis

Data are presented as the mean ± SD. Comparisons were conducted using Student’s t test, assuming a two-tailed distribution, or by one-way ANOVA with Tukey’s post test, with significance defined as P < 0.05 (*), P < 0.01 (**), or P < 0.001 (***). The number of biological replicates for each experiment is indicated in the corresponding figure legend. Histological and immunofluorescence images are representative of at least 3 different experiments/animals.

---

**Suppl. Figure 1:**
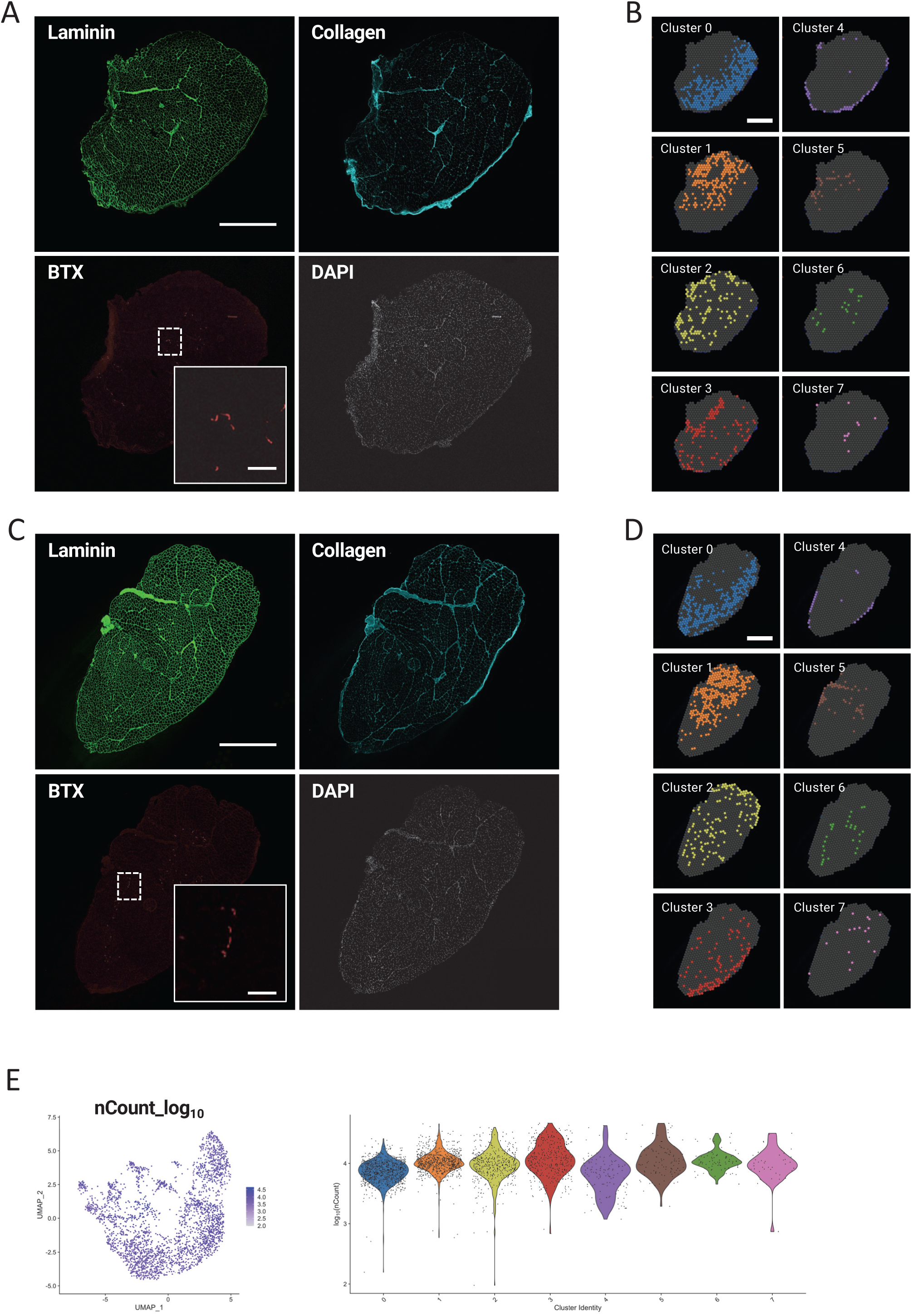
A) 3 days post nerve crush mouse hindlimb cryosection used of ST. Immunofluorescence staining for Laminin (top left) Collagen (top right) Bungarotoxin (bottom left) Hoechst (bottom right). Scalebar 1mm B) Spatial distribution of each cluster from ST data in A. Scalebar 1mm, Inset Scale Bar 100um. 30 days post nerve crush mouse hindlimb cryosection used of ST. Immunofluorescence staining for C) Laminin (top left) Collagen (top right) Bungarotoxin (bottom left) Hoechst (bottom right). Scalebar 1mm, Inset Scale Bar 100um. D) Spatial distribution of each cluster from ST data in C. Scalebar 1mm E) Log10 UMI distribution plotted over UMAP embedding (left) or divided per cluster (right) displaying a comparable number of unique transcripts in the integrated dataset and in the different clusters

**Suppl. Figure 2:**
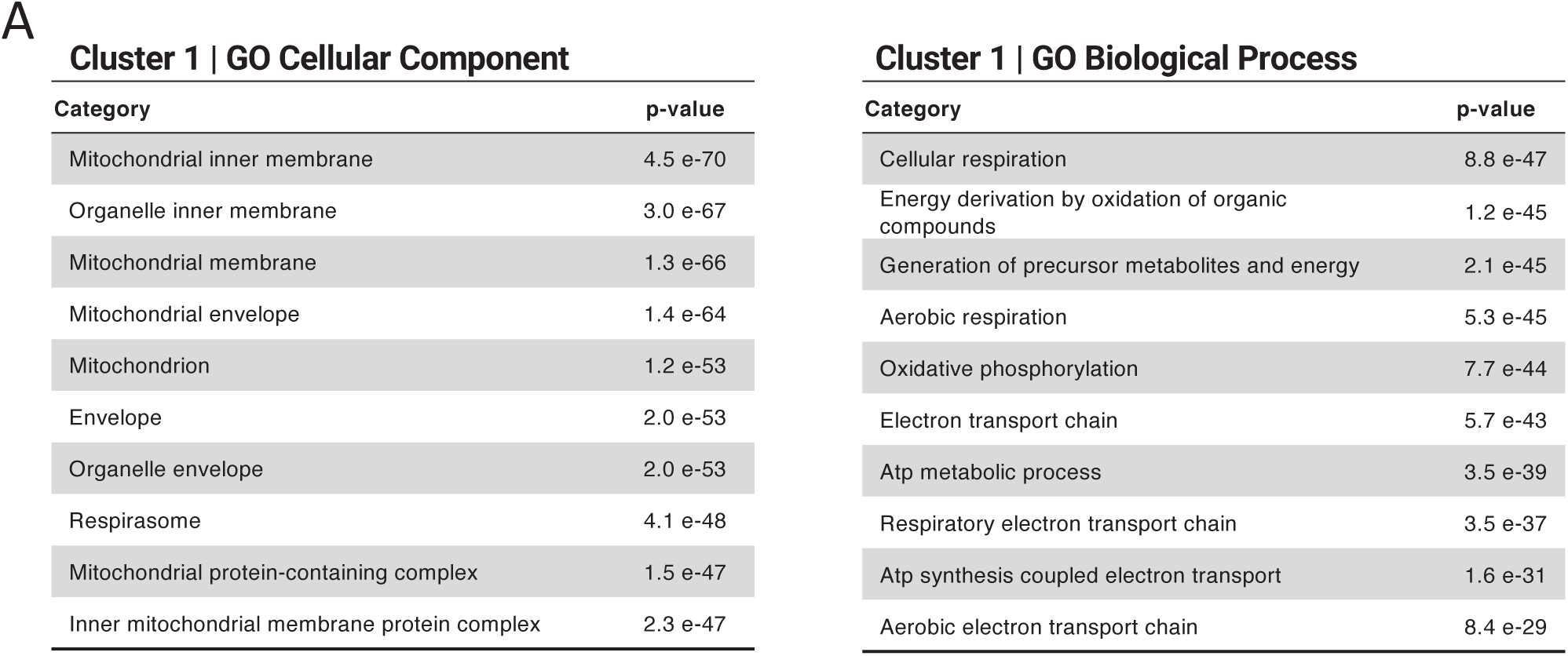
A) Cellular Component and Biological Process Gene Ontology analyses of Cluster 1

**Suppl. Figure 3:**
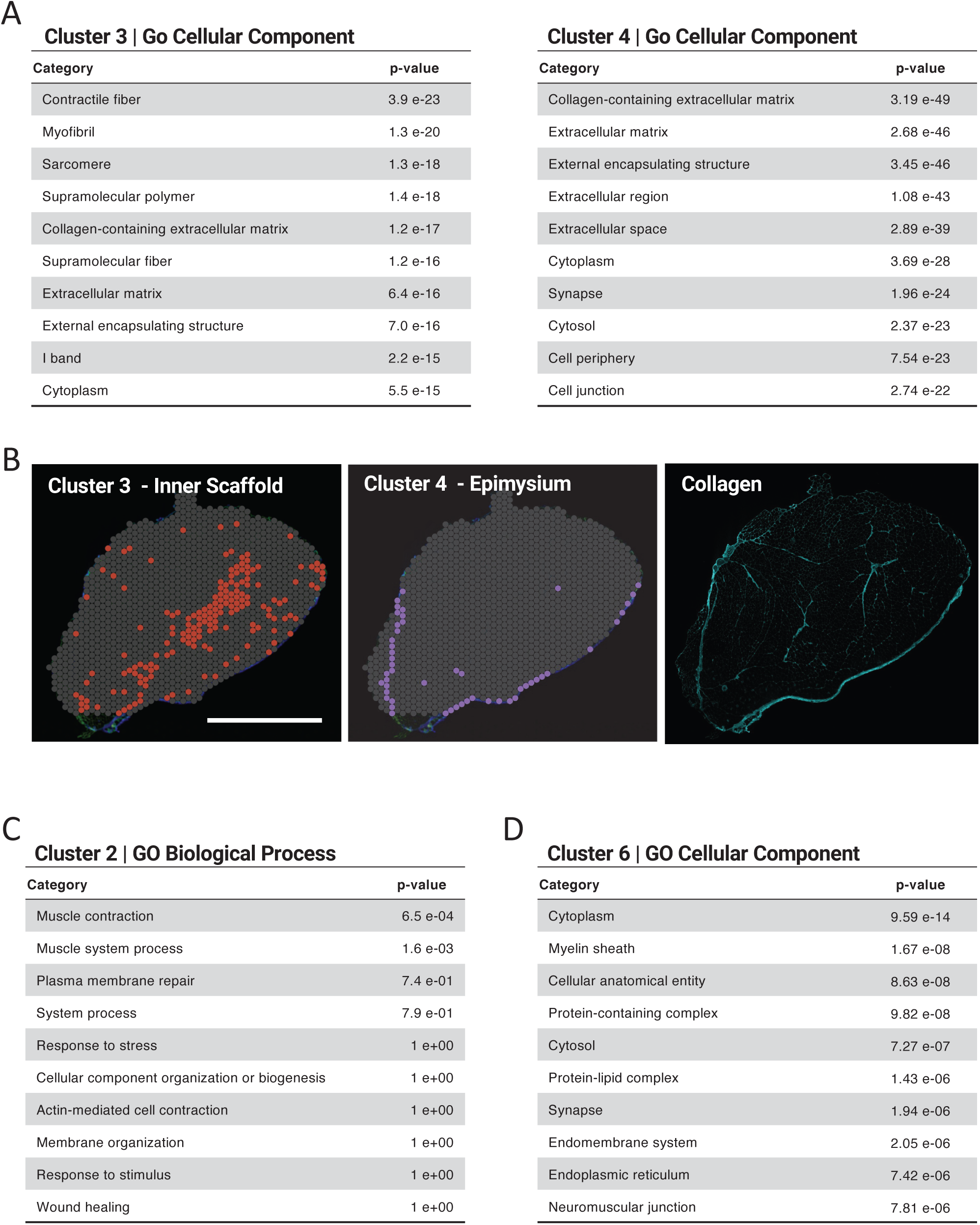
A) Cellular Component Gene Ontology enrichment analysis of Cluster 3 and 4 B) Spatial distribution of cluster 3(left) and Cluster 4 (middle). Immunofluorescent analysis of Collagen 1 (Right). Scalebar 2mm C) Biological Process Gene Ontology analysis of Cluster 2 D) Cellular Component Gene Ontology enrichment analysis of Cluster 3

**Suppl. Figure 4:**
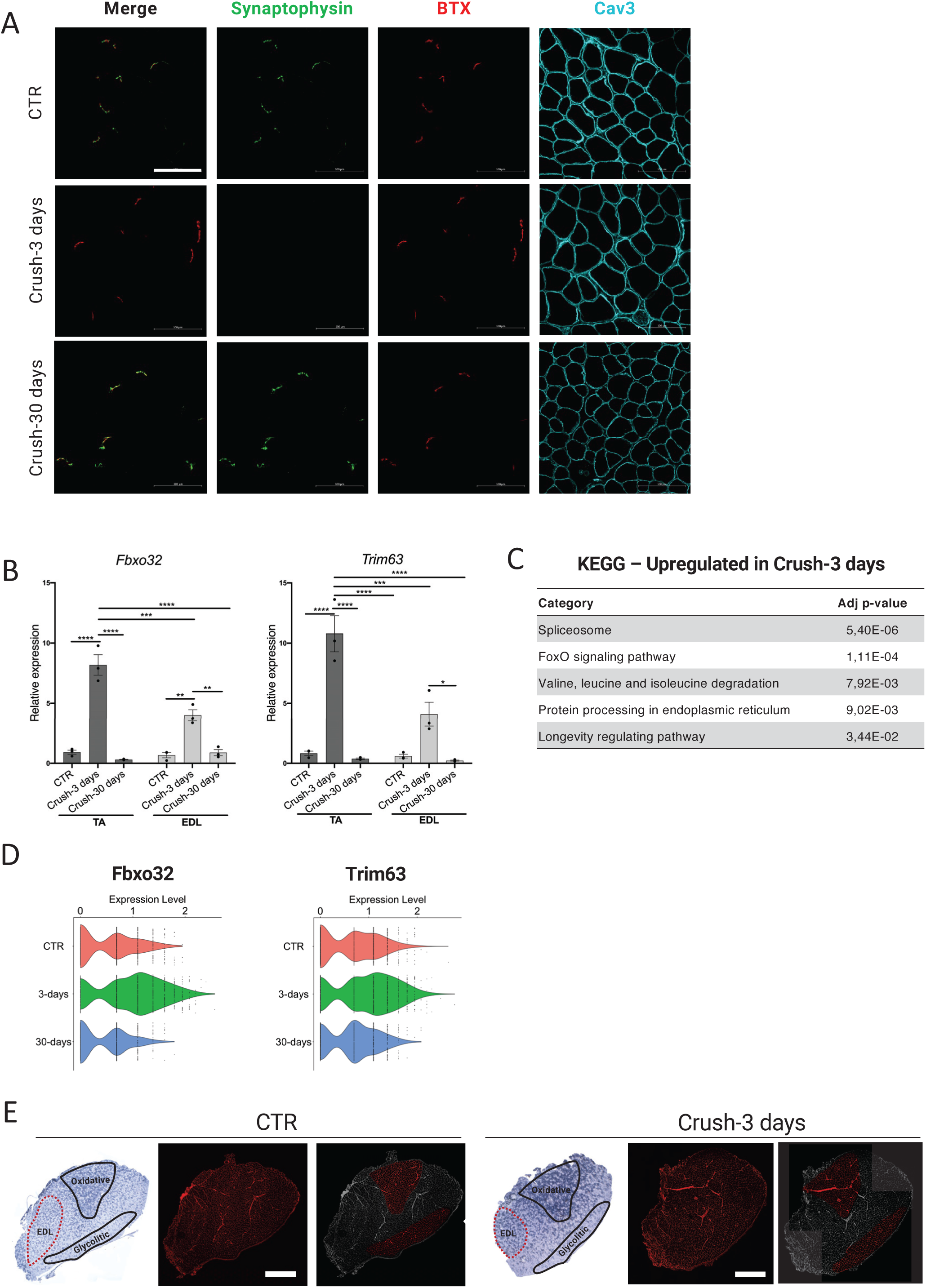
A) Representative immunostaining for Synaptophysin (green), bungarotoxin (red) and Caveolin-3 (cyan), the first image is an overlay among synaptophysin and bungarotoxin. Scale 0.5 mm B) Relative expression of Fbox32 Trim63 during denervation in TA and EDL muscle (n = 3, values represent mean SD, ***P < 0.001, ****P < 0.0001; by 1-way ANOVA Tukey’s multiple-comparison test). C) Pseudo-bulk KEGG pathway enrichment analysis of whole Crush 3 days sample. D) Violin plot showing the expression of Fbox32 Trim63 in the different ST timepoints. E) Muscle section compartmentalization strategy for microdissection. Immunostaining for Laminin (red). Scale 1 mm Areas highlighted in red have been used to calculate CSA from oxidative and glycolytic parts of the muscle

**Suppl. Figure 5:**
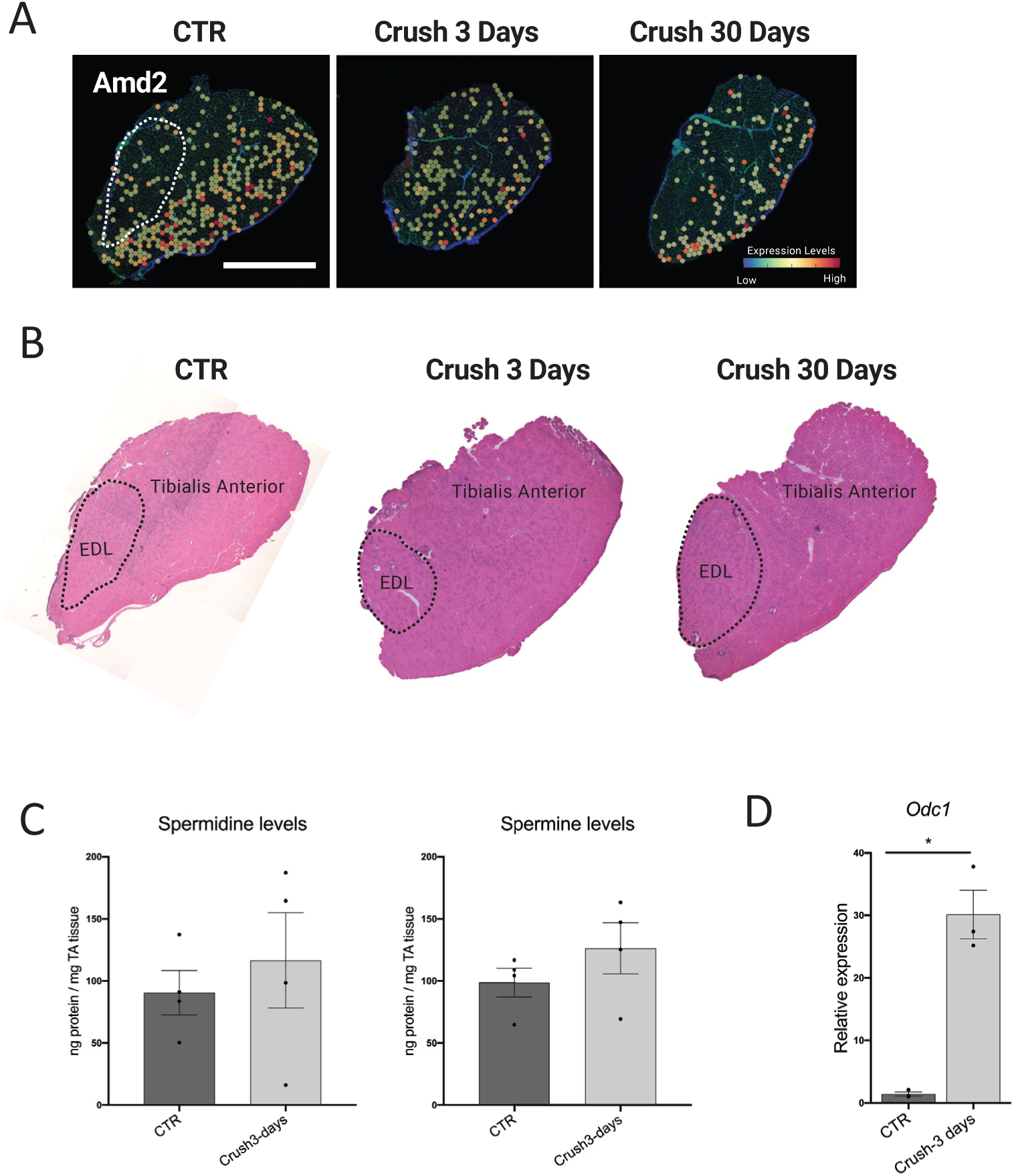
A) Relative expression levels of Amd2 scale 2mm (white dotted lines highlights EDL muscle) B) Serial section stained with Hematoxylin and Eosin illustrating muscle localization C) GC-MS quantitative analysis of Spermidine and Spermine levels in control (CTR) and denervated (Crush 3 days) muscles (n = 4, values represent mean SD by 2-tailed Student’s t-student test). D) Relative expression of Odc1 during denervation in TA (n = 3, values represent mean SD,*P < 0.05 by 2-tailed Student’s t-student test).

